# Sea surface freshening can suppress the thermal tipping point of marine copepods

**DOI:** 10.1101/2025.09.23.677975

**Authors:** Mathieu Lutier, Andrea Emilie Thorstensen Skari, Helena Reinardy, Heidi Sjursen Konestabo, Katrine Borgå, Khuong V. Dinh

**Affiliations:** Section for Aquatic Biology and Toxicology, Department of Biosciences, University of Oslo, Oslo, Norway; Scottish Association for Marine Science (SAMS), The University of the Highlands and Islands (UHI), Oban, Scotland, United Kingdom; Department of Arctic Biology, The University Centre in Svalbard (UNIS), P.O. Box 156, Longyearbyen, Norway; The Science Library, University of Oslo, Oslo, Norway

**Author notes:** Co-first authors (Mathieu Lutier, Andrea Emilie Thorstensen Skari). Corresponding authors: Mathieu Lutier and Khuong V. Dinh.

**Keywords:** *Calanus* spp., Degree day, Marine heatwaves, Performance curve, Reaction norm, Salinity, Warming

## Abstract

Tipping points govern species distributions, which could be impacted by ocean warming (OW) and sea surface freshening (SSF). Interactions between these stressors may affect tipping points. Also, the accumulation of stress over time (degree day) could condition the reaching of tipping points. Yet, these aspects have never been studied. Here, we hypothesise that SSF reduces the thermal tipping point of *Calanus* spp., key copepods that play a structuring role in marine food webs. *Calanus* spp. from the Oslofjord were exposed to a temperature gradient from 8 to 20 °C and a salinity range from 35 to 10 PSU. We found that the survival of *Calanus* spp. tipped and decreased above temperatures of 14 – 18 °C, which are already reached in the surface waters of the Oslofjord during summer months. At 29 PSU, this corresponded to a tipping point of heat accumulation of 28 °C.d above which survival decreased. Reduced salinity below 29 PSU, prevalent year-round in surface waters, suppressed this tipping point, inducing linear decreases in survival when temperature rises above 8 °C. Thus, the thermal niche of *Calanus* spp. will decrease with SSF, impacting whole ecosystems. We provide the first statistical determination of heat accumulation tipping points. We also demonstrate that exposure to a secondary stressor can suppress tipping points to a primary stressor, thereby removing the ability of phenotypic plasticity to buffer negative impacts on fitness. This advances our knowledge of physiological tipping points, which are essential in a world where they are increasingly reached by environmental changes.

## Introduction

Climate change is driving rapid ocean warming (OW), with global ocean surface temperatures projected to increase by 1 to 5°C by 2100 [1,2]. In high latitudes, increased precipitation is reducing sea surface salinity, causing sea surface freshening (SSF) [3]. Concurrently, climate change is also increasing the frequency, intensity, and duration of extreme weather events, causing extreme and sudden warming, i.e., marine heatwaves (MHW) and salinity drops [4]. Such extreme events of warming, with temperatures up to 5°C higher and salinity drops up to 15 PSU lower than seasonal norms, can last from several days to months [5–7]. These rapid temperature and salinity changes affect species survival, physiology, metabolism, reproduction, behaviour, and phenology [8–10]. As a defence, organisms rely on phenotypic plasticity by adjusting their physiology, metabolism, and behaviour to maintain their performance, primarily their survival and reproduction [11].

Phenotypic plasticity has limits, and exceeding tolerance thresholds can cause irreversible damage, from molecules to the entire organism [12,13]. The tolerance threshold to OW is generally studied as the Critical Thermal maximum (CTmax), which is the temperature above which there is a loss of function (e.g. cessation of movement, cessation of growth, death) [14] (Figure 1). In contrast, the tipping point is the threshold, e.g., a temperature value, beyond which stress begins to have a brutal, sudden, and irreversible effect on physiology, i.e. homeostasis is compromised [15–17], and ultimately on fitness, i.e. survival and reproduction [12,13] (Figure 1). Beyond tipping points, populations have to adapt over long-term, evolutionary timescales or face extirpation [12,13]. Crossing the tipping points of keystone species, e.g., providing food or habitats to other species, could ultimately lead to the reaching of ecosystem tipping points [18–21]. Beyond tipping points, ecosystems transition from one stage to another, affecting the services they provide to human societies.

**Figure 1.**
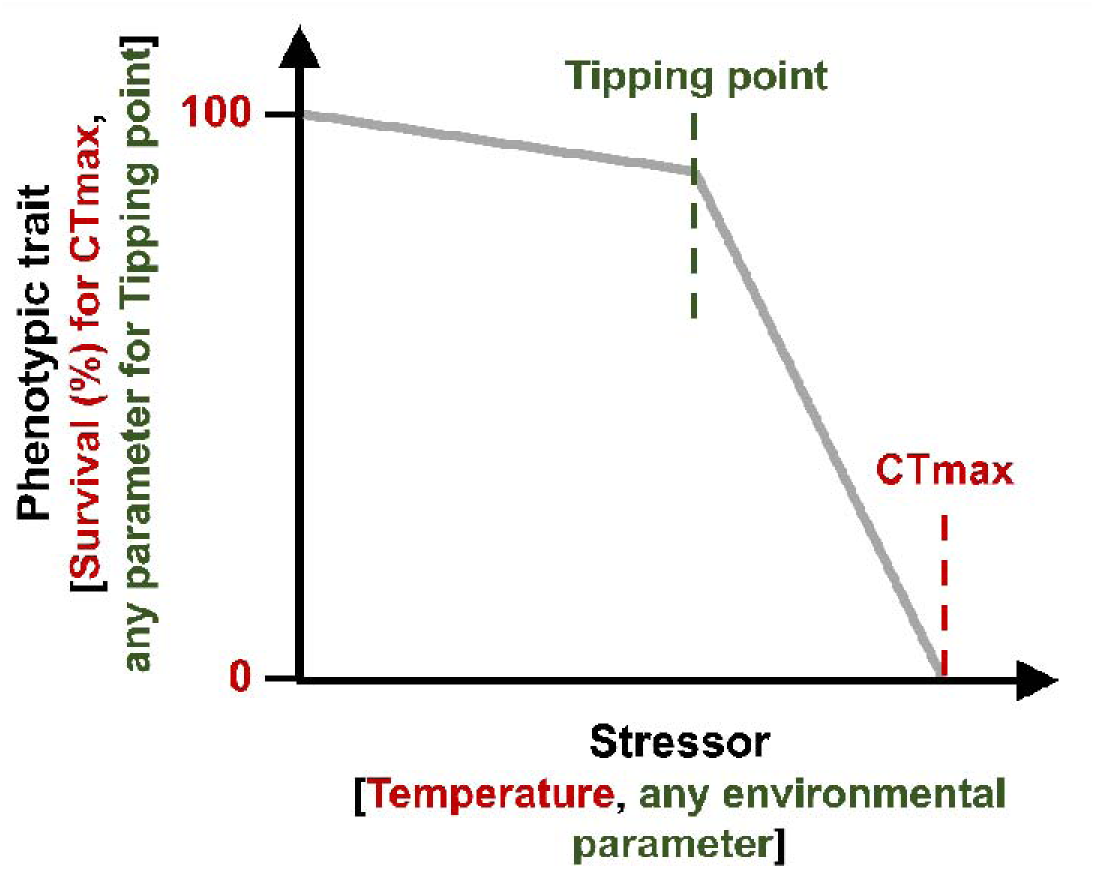
Example of a performance curve (note that shape can be very different, e.g. no tipping point), showing different kinds of stress tolerance thresholds: tipping point (green) and critical thermal maximum (CTmax, red). Tipping points can be determined for any type of physiological traits and stressors. CTmax can only be determined for temperature.

Determining species tipping points involves determining performance curves, which are the modelling of biological parameters across a wide range of environmental conditions. This is experimentally challenging and requires significant resources. Consequently, performance curves are rarely studied for multiple stressors, complicating projections about species’ future [16,22]. Stressor interactions can result in the tipping point for one stressor to be reached earlier when exposed to an additional stressor [20,23]. SSF has been shown to decrease the tolerance of several marine organisms to OW [24], and could therefore decrease their thermal tipping points. Yet, this has not been studied.

In addition to stress intensity and multiple stressor interactions, exposure duration is another crucial parameter for predicting the impact of global changes on the physiology of organisms [25]. For example, it has been shown that coral death only occurs after a certain time spent above a temperature threshold [26]. Therefore, determining accumulated heat stress, calculated by integrating temperature over time (degree days, degree hours, etc.), is essential for predicting the negative effects of MHW on organisms [25–28]. It is therefore likely that there is a tipping point in heat accumulation (degree time) beyond which the fitness of marine organisms declines. However, this time-sensitive aspect of tipping points remains unknown.

Tolerance thresholds, such as tipping points, to temperature and salinity govern species distributions, which are increasingly altered by climate change [29,30]. A striking example is the species of the North Sea copepod community, whose distribution is projected to shift northwards at a rate of 9 km per decade because of OW [31]. We hypothesize that this negative impact of OW could be amplified by strong SSF in the North Sea, that would decrease the thermal tipping point of copepods. This could be particularly the case in the Skagerrak, at the mouth of the Baltic Sea, where SSF is among the fastest on the planet [32]. OW and SSF in interaction have been shown to negatively impact the survival and metabolism of *Calanus* spp. [33]. *Calanus* spp. are particularly lipid-rich and generally dominate zooplankton biomass in North Atlantic ecosystems [34,35]. They thus play a key structuring role in marine food webs [36], and any changes impacting their distribution would affect entire ecosystems.

Here, we test the hypothesis that SSF decreases the thermal tipping point of *Calanus* spp.. We exposed *Calanus* spp. collected in the Oslofjord (North Skagerrak) to temperatures from 8 to 20 °C and salinity from 35 to 10 PSU. During the exposure, temperature was recorded at high frequency and integrated as a function of time to determine heat accumulation (°C.d). Survival was recorded for 8 days to determine the heat accumulation tipping point and how it varied as a function of SSF. It has recently been shown that sublethal effects can affect organisms even before reaching tipping points [17,37,38]. Thus, to detect sublethal effects, we also measured the amount of DNA damage, i.e., genotoxicity, that can reflect oxidative stress caused by heat shock [39]. Gene expression is the basis for higher-level physiological responses, and, therefore, DNA damage can have cascading effects on the survival capacity of organisms [40,41]. Here, we assess how structural genotoxicity mechanisms might be linked to reaching survival tipping points. Warming tipping point decreasing with SSF would have strong consequences for the future distribution of species. This study is particularly relevant in a world where tipping points are increasingly crossed by environmental changes but are never studied under the interactive effects of multiple stressors and duration of exposure.

## Materials and methods

### Sampling

Copepods were collected in the Oslofjord, on 9 October 2023, near IM2 Station (59°37’ N, 10°37’E), onboard the research vessel *F/F Trygve Braarud*. Sampling was conducted by vertically hauling a WP2 net (mesh size 200 μm) from 198 to 40 m depth. The seawater temperature and salinity below the thermocline, where most calanoid copepods are distributed during the day [42,43], were recorded with a CTD and were approximately 8 °C and 32 PSU. Live samples were diluted in seawater collected in the deep, stored in coolers, and transported to the University of Oslo (UiO). Sorting was conducted in a climate room at ∼8 °C using a stereomicroscope. Samples were separated by species and life stages based on morphological criteria [44]. We identified *Calanus* spp. at the copepodite CV stage as the dominant group in the community, which was selected for the experiment. Here, the *Calanus* spp. were assumed to be mostly *Calanus helgolandicus* (Claus, 1863) based on previous observations [45,46], but could be a mix with the morphologically similar *Calanus finmarchicus* (Gunnerus, 1770). Sorted copepods were kept in open bottles of filtered seawater at a density of 45 copepods/L in an incubator at 8 °C. The same batch of seawater was used to maintain copepods throughout the experiment. This seawater was collected at Drøbak Aquarium (pumped at 40 m depth in the Oslofjord). The seawater was filtered at 1 μm, treated with a UV lamp, and stored at 4 °C. Air was bubbled into the seawater for good oxygenation.

### Experimental design

Copepodite stage CV *Calanus* spp. were exposed to 5 temperatures, 8, 11, 14, 17, and 20 °C, and 5 salinities, 34, 28, 22, 16, and 10 PSU. These conditions cover the range of temperatures and salinities recorded on the whole water column at the IM2 sampling station during monthly measurements in 2022 that ranged from 3 to 20 °C and from 35 to 22 PSU. Extreme conditions (20 °C and salinities < 22 PSU) were added to challenge organisms’ physiology and detect tipping points, as recommended in the literature [17,47]. Each temperature condition was maintained in one of 5 identical incubators. Salinity treatments were prepared by mixing distilled water with seawater and measuring salinity with a conductivity probe (Multiline ® Multi 3630 IDS-WTW, TetraCon® 925 WTW).

On 18 October 2023, individual copepods were transferred in plastic flasks (VWR Tissue culture flasks, surface treated, sterile, vented caps, 25 cm^2^) filled with 65 mL of seawater at the different salinities. 15 copepods were exposed for each salinity treatment. Then, 75 copepods, corresponding to the 5 salinity treatments, were placed in each of the 5 incubators set at an initial temperature of 8°C. For 4 days, the temperature was then increased sequentially by 3 °C.d^-1^ so that all temperature conditions were reached on the same day in the different incubators (Supplementary Figure 1). The exposure to constant temperatures then lasted 8 days, a common duration for MHW in nature [5]. Both *C. finmarchicus* and *C. helgolandicus* have large lipid reserves to survive the low-feeding season that last several months in the North Sea [48–50]. Therefore, these lipid reserves are unlikely to have decreased during the short duration of the experiment and to have affected survival.

### Daily measurements

Survival was checked daily by tapping the wall of the flask or gently touching the copepods with a pin to stimulate them. Immobile copepods were considered dead. Salinity was measured daily in the flasks of the dead copepods. On the final day, salinity was measured in three flasks from each condition. These measurements aimed to ensure that evaporation did not change salinities over the course of the experiment. The average salinity in each condition was then calculated and used for statistics. Temperature was recorded in each incubator every 15 minutes using HOBO^©^ TidbiT v2 loggers (1 logger/incubator) placed in a 65 ml beaker, sealed with parafilm, and filled with seawater. The average temperature in each incubator during the exposure period was calculated and used for statistics.

We also calculated the heat accumulation from the time the copepods were transferred into the flasks (Supplementary Figure 2), as follows:

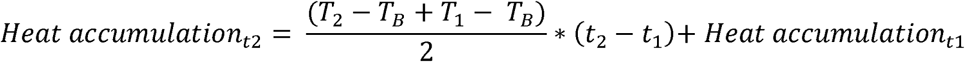

*Heat accumulation_t1_* and *Heat accumulation_t2_* are the heat accumulations (°C.d) at timepoints *t1* and *t2* (d) respectively. *T_1_* and *T_2_* are the temperature (°C) at *t_1_* and *t_2_*. This corresponds to the calculation of the area under the temperature-time curve and above the base temperature, *T_B_*, of 8 °C (control temperature in the Oslofjord at the time of sampling). This is done by estimating the area of a trapezoid between two temperature measurements [51]. This method is commonly used to assess the influence of MHW on coral bleaching [25–28]. As an example, in our case, 0, 1 and 2 °C.d could represent 0, 1, and 2 days spent at 9 °C, i.e., +1°C above the control temperature of 8 °C, respectively. Alternatively, 2°C.d could, for example, represent 1 day spent at 10°C or 4 days at 8.5°C.

### DNA damage analyses

The fast micromethod assay (FMM) was used for determining the amount of DNA damage by assessing the unwinding kinetics of double-stranded DNA in alkaline solution with fluorescence measurements [52,53]. For each temperature/salinity condition, at the end of the experiment, 5 ± 2 copepods (depending on the final survival, Supplementary Table 1) were individually snap**-**frozen in liquid nitrogen and stored at −80 °C. Our protocol is adapted from Halsband *et al.* [54]. Individual copepods were gently homogenised with a hand pestle in 260 μL of 20 mM EDTA + 10% DMSO. 20 μL of homogenate, or 20 mM EDTA + 10% DMSO for the blank, was loaded, with 5 analytic replicates per sample, into a microplate (96-well black-walled, Greiner Bio-One Ltd). Samples were lysed on ice for 40 min in the dark after adding 20 μL of lysis buffer to each well (20µl 9M urea, 0.1% SDS, 0.2M EDTA, 2% Quant-iT™ PicoGreen®). After lysis, unwinding was initiated by adding 200 μL of unwinding solution (mixture of 20 mM EDTA and 2 M NaOH until pH is adjusted to 13.0) to each well. Fluorescence was then recorded immediately using a SynergyMx^®^ microplate reader (BioTek^©^) and every minute for 30 min (excitation 485nm, emission 520nm). The amount of DNA damage was calculated as the strand scission factor [53], as follows:

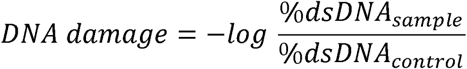

*%dsDNAsample* is the percentage of double-stranded DNA in each sample and *%dsDNAcontrol* is the average percentage of double strand DNA in the controls, i.e. the copepods exposed at 8°C. The *%dsDNA* values were calculated as the fluorescence (RFU) measured for a sample at 20 min, divided by the average RFU recorded in the controls at 0 min.

### Statistical analyses

All analyses were performed using the R software v4.3.0, and the statistical significance threshold was 0.05. The effects on survival of temperature, salinity and the interaction of salinity and temperature, were tested using Cox proportional-hazards model using the *coxph* function of the *survival* package [55]. The assumption of proportional hazards of the Cox model was graphically checked by plotting the Schoenfeld residuals against time using the *ggcoxzph* function of the *survminer* package [56]. For significant factors, i.e., temperature salinity and/or their interaction, *post-hoc* pairwise comparisons were performed using the *pairwise_survdiff* function (using the Benjamini-Hochberg procedure). We did not include DNA damage as a factor influencing survival because it is not measurable in dead copepods.

For each salinity, we aimed to determine the thermal tipping point, in °C, which was not possible by modelling survival as a function of temperature. Indeed, we had only 5 data points of survival as a function of temperature, which is not enough for piecewise regressions [57]. However we determined the heat accumulation tipping point, in °C.d, by modelling daily survival (%) as a function of the heat accumulation, since the start of the experiment, calculated from high-frequency temperature measurements (see above). We thus had 60 data points to model performance curves, by pooling daily survival data recorded in each of the 5 incubators, for each salinity condition (Supplementary Figure 2). We tested piecewise, linear, and log-linear regressions to model survival against heat accumulation, choosing the best model according to statistical criteria, as described in [58]. Piecewise regression detects slope breaks in the regression corresponding to a tipping point, the factor value (i.e. heat accumulation) where the variable (i.e. survival) tips, using the bootstrap restarting algorithm (Figure 1) [57]. Slopes and tipping points were then compared between each salinity condition using Welch t-test and the 95% confidence intervals of the parameters given by the models, as described in [38]. All slopes and tipping points were statistically different between the different models across the different salinity conditions,

Finally, type II ANOVA, for unbalanced designs (Supplementary Table 1), was used to test the effects of temperature, salinity, and their interactions on DNA damage. The homogeneity of variances and normality of residuals were checked graphically. The differences between groups for significant factors were tested using Tukey HSD tests.

## Results

Temperature and salinity were stable during the 8 days of exposure and were 8.3 ± 0.0, 11.0 ± 0.0, 13.9 ± 0.0, 17.7 ± 0.1, and 20.2 ± 0.1 °C, and 32.2 ± 0.2, 28.5 ± 0.2, 22.5 ± 0.2, 16.7 ± 0.1, and 10.3 ± 0.1 PSU, respectively. We found a significant effect of temperature (*p < 0.001*), salinity (*p < 0.001*), and the interaction of temperature and salinity (*p < 0.001*) on copepod’s survival (Figure 2). Survival was globally similar between 8-14 °C, while survival decreased above 14-18 °C (global post-hoc test for temperature). Survival was globally similar between salinities 32-23 PSU (global post-hoc test for salinity). Survival curves were significantly different at 17 PSU, with low survival after 8 days, and 10 PSU, with sudden death as soon as the copepods were transferred into the flasks. This suggests tipping points for temperature and salinity, at 14-18°C and 23-17 PSU, beyond which survival is strongly affected.

**Figure 2.**
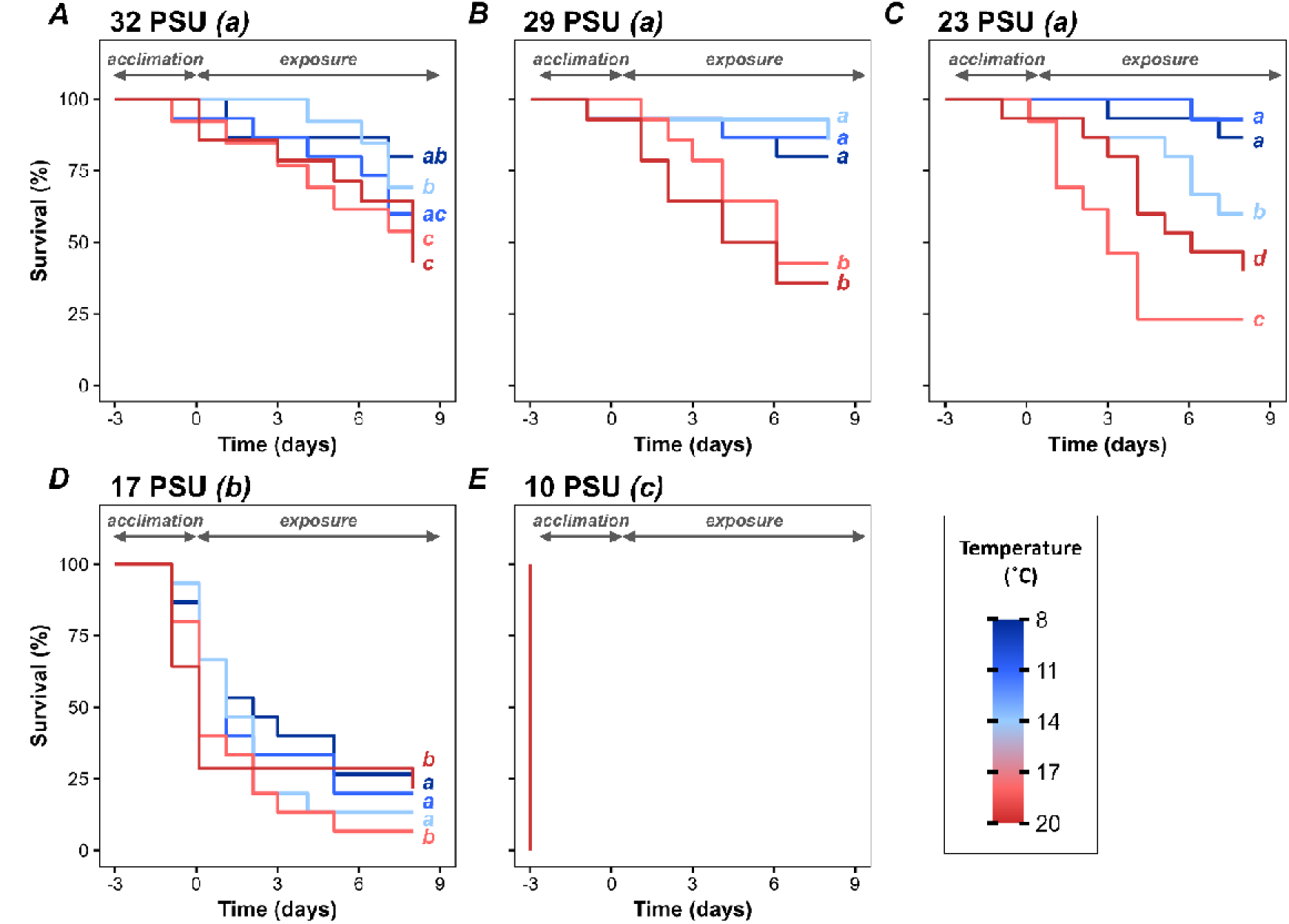
Kaplan-Meier survival curves showing survival (%) of CV Calanus spp. over time, as a function of temperatures (8 – 20 °C) and different salinities (32 – 10 PSU). Salinities are A) 32 PSU, B) 29 PSU, C) 23 PSU, D) 17 PSU, and E) 10 PSU. The temperature during the exposure period is represented by the colour gradient. Survival curves that differ significantly for different salinities and temperatures are indicated by different letters in the title or the figure, respectively. Negative values for time on the x-axis indicate the days before the designated experimental temperature condition is reached, i.e., the “acclimatisation” period. The exposure period is indicated by positive values.

The significant interaction between salinity and temperature showed that survival curves reacted differently to warming depending on the salinity. At 32 PSU, survival decreased significantly with warming, overall in a linear manner (post-hoc tests, Figure 2A). At 29 PSU, there was no difference in survival until reaching a tipping point at 14-18 °C, above which survival declined sharply (post-hoc tests, Figure 2B). At 23 PSU, there was no difference in survival until reaching a tipping point at 11-14 °C, above which survival decreased sharply (post-hoc tests, Figure 2C). At 17 PSU, survival decreased over time with almost no notable difference between temperature conditions (post-hoc, Figure 2D). At 10 PSU, copepods died from salinity shock before temperature had any effect (Figure 2E).

The performance curves of survival as a function of heat accumulation (°C.d) provided further insight into the effects of salinity on thermal tolerance (Figure 3). At 32 PSU, survival decreased linearly as soon as heat accumulated, at a rate of -0.4 % °C.d^-1^ (Figure 3A). At 29 PSU survival is not affected (*p > 0.05*), until reaching a tipping point at 28 °C.d, beyond which survival decreased linearly with heat accumulation at a rate of -0.7 % °C.d^-1^ (Figure 3B). At 23 PSU, the tipping point disappeared and, survival decreased linearly as soon as heat accumulated, at the rate of -0.7 % °C.d^-1^ (similar pattern but much faster than at 32 PSU) (Figure 3C). At 17 PSU, the pattern reversed and survival declined extremely rapidly as heat accumulated, at a rate of -32% °C.d^-1^, before reaching a tipping point at 2 °C.d, beyond which it declined more slightly (-0.3% °C.d^-1^) (Figure 3D). At 10 PSU, copepods died before warming had any effect on survival. Importantly, survival is not affected by heat accumulation (i.e. time spent above 8 °C) only at the salinity of 29 PSU, before reaching the tipping point at 28 °C.d (Figure 3B). Reaching this tipping point would be equivalent to spending 28 days at 9°C or 4 days at 15°C, for example. With increasing salinity, 32 PSU (Figure 3A), and decreasing salinity, 23 PSU (Figure 3C), vulnerability to warming increases since the tipping point disappears and survival is affected as soon as heat accumulates. The vulnerability to warming is extreme at 17 PSU with survival dropping from 100 % to 37 % between 0 and 2 °C.d (Figure 3D). 2 °C.d is equivalent to spending two days at 9 °C.d. The tipping point detected by the piecewise regression at 17 PSU is a statistical tipping point, i.e, a break in the slopes, but not a physiological tipping point. Indeed, most individuals died as soon as heat accumulated and the decrease in survival is anecdotal after 2 °C.d, because most copepods were already dead. In contrast, at 29 PSU, there was no significant decrease in survival before reaching the physiological tipping point at 28 °C.d.

**Figure 3.**
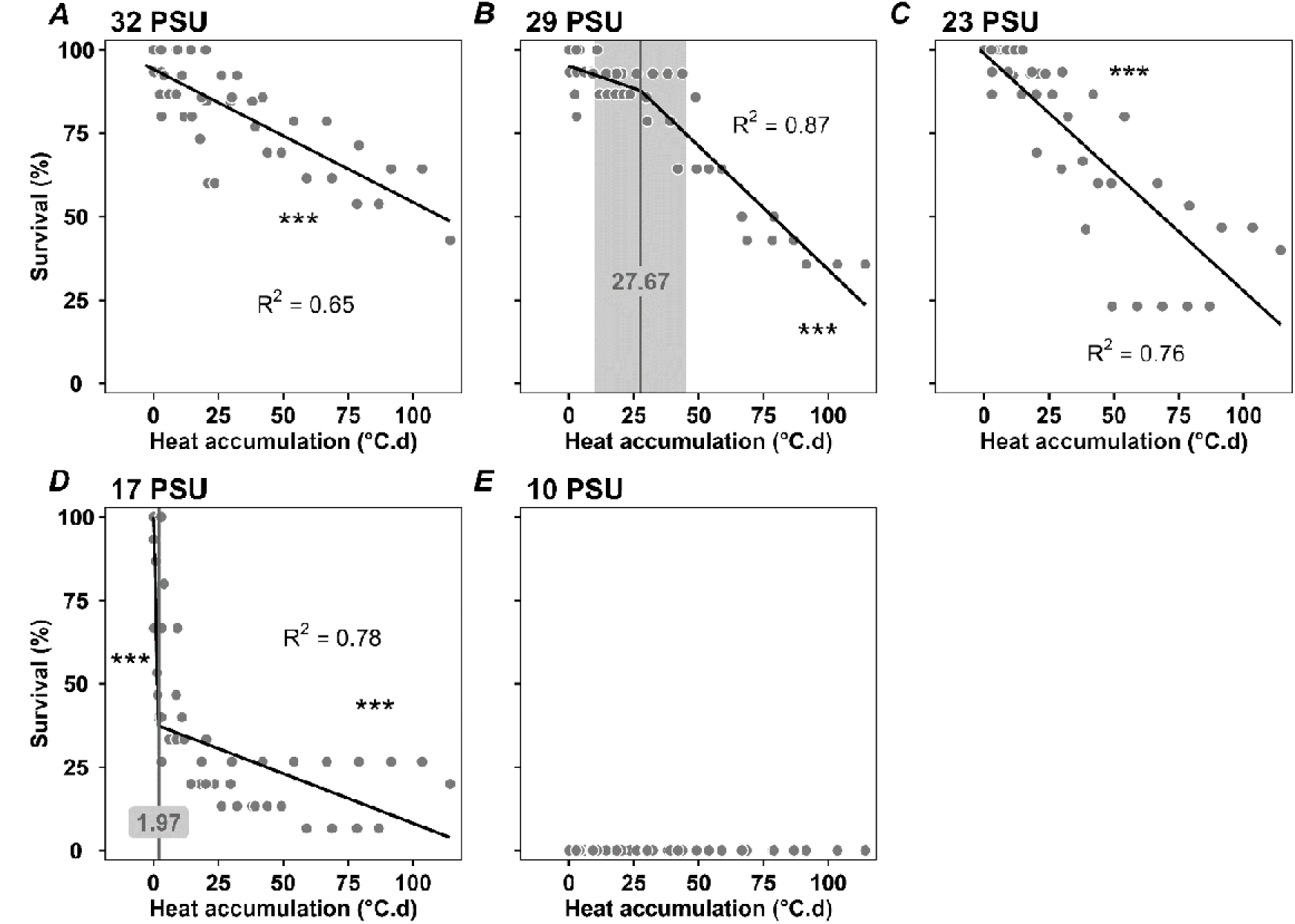
Survival of CV *Calanus* spp. (%) as a function of heat accumulation since the beginning of the experiment (°C.d) at different salinities (32 – 10 PSU). Salinities are A) 32 PSU, B) 29 PSU, C) 23 PSU, D) 17 PSU, and E) 10 PSU. Heat accumulation represents the time spent above 8 °C, with 1 °C.d representing, for instance, +1 °C (9 °C) for 1 d or +0.5 °C for 2 d. Survival was modelled with linear or piecewise regressions. Tipping points and their 95% confidence intervals are shown in grey. The significance levels of the slopes are presented using symbols (p < 0.001 ***, p <0.01 **, p < 0.05 *). In the absence of a symbol, a slope is not significantly different from 0.

The interaction of temperature and salinity had no effect on DNA damage (*p = 0.714*). Temperature had no effect on DNA damage (*p = 0.366*, Figure 4A), which was only significantly affected by salinity (*p < 0.001*, Figure 4B). There was no difference in DNA damage between 32 and 23 PSU, with an increase below a tipping point at 23-17 PSU. This was consistent with the tipping point observed for survival when salinity decreased according to the Cox Proportional-Hazards Model (Figure 2).

**Figure 4.**
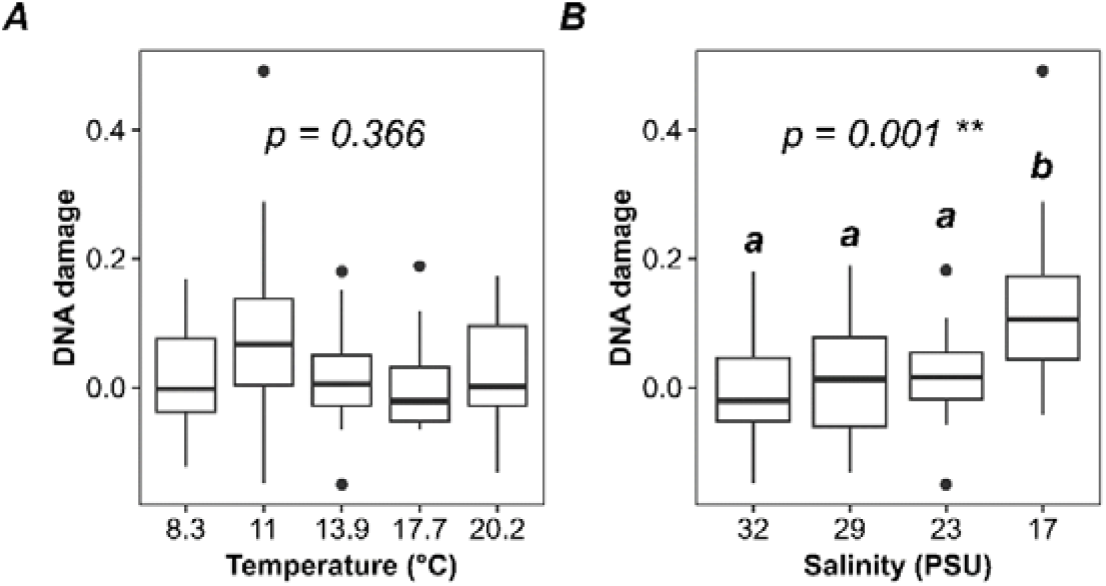
DNA damage in CV *Calanus* spp. that survived 8 days of exposure to warming, at 5 different salinities. Data are presented as box and whisker plots (averages and standard deviations) as a function of A) temperature and B) salinity, across all salinities and temperatures, respectively. The p.value of type II ANOVA is displayed for each factor. The results of Tukey HSD post-hoc test are shown with letters, for salinity (significant). DNA damage that differs significantly is indicated by different letters.

## Discussion

Tolerance thresholds, such as tipping points to temperature and salinity, are among key factors determining the distribution of marine species, which are impacted by global changes. Multiple environmental changes are occurring at the same time, and exposure to one additional stressor could shift the tipping point to another stressor. Yet the influence of multiple stressors on tipping points is rarely studied. Here, we hypothesize that exposure to SSF would accelerate the reaching of a warming tipping point in *Calanus* spp.. These species are key copepods in pelagic ecosystems of the North Seas and Skagerrak [34–36], where OW and SSF are among the fastest recorded on the planet [32]. We also assess, for the first time, whether the accumulation of heat stress over time (°C.d) leads to reaching a tipping point in the survival of marine organisms.

We show that survival of *Calanus* spp. is impacted above 14-18 °C (cox model). Our results could be affected by the presence of two species *C. finmarchicus* and *C. helgolandicus* in our experiments that have different tolerances to warming, and are both present in the Skagerrak [59,60]. *C. finmarchicus* is a boreal species that is generally abundant at 0-9°C, while *C. helgolandicus* is more temperate and abundant at 13-18 °C [34]. *C. finmarchicus* survival has been shown to decrease above 15°C with no survival above 20°C [61], corresponding to our results. The thermal tolerance of *C. helgolandicus* has not been assessed, but is expected to be higher [34]. We show that the survival of *Calanus* spp. is impacted below a salinity tipping point at 23-17 PSU, with no survival at 10 PSU. This is similar to what has been previously observed on *C. finmarchicus* from Disko Bay, Greenland [33]. *C. finmarchicus* and *C. helgolandicus* have generally maximum abundances at salinity of 34-35 and 35-36 PSU, respectively, with *C. helgolandicus* being more tolerant [59]. Surface temperatures in the Oslofjord in 2022 already exceeded the 14-18 °C tipping point for 4 months from June to September (Figure 5A). Surface salinity only crossed the 23-17 PSU tipping point for survival for a few days in May 2022 (Figure 5B). Therefore, the survival of *Calanus* spp. may already be compromised in the surface waters of the Oslofjord for short periods.

**Figure 5.**
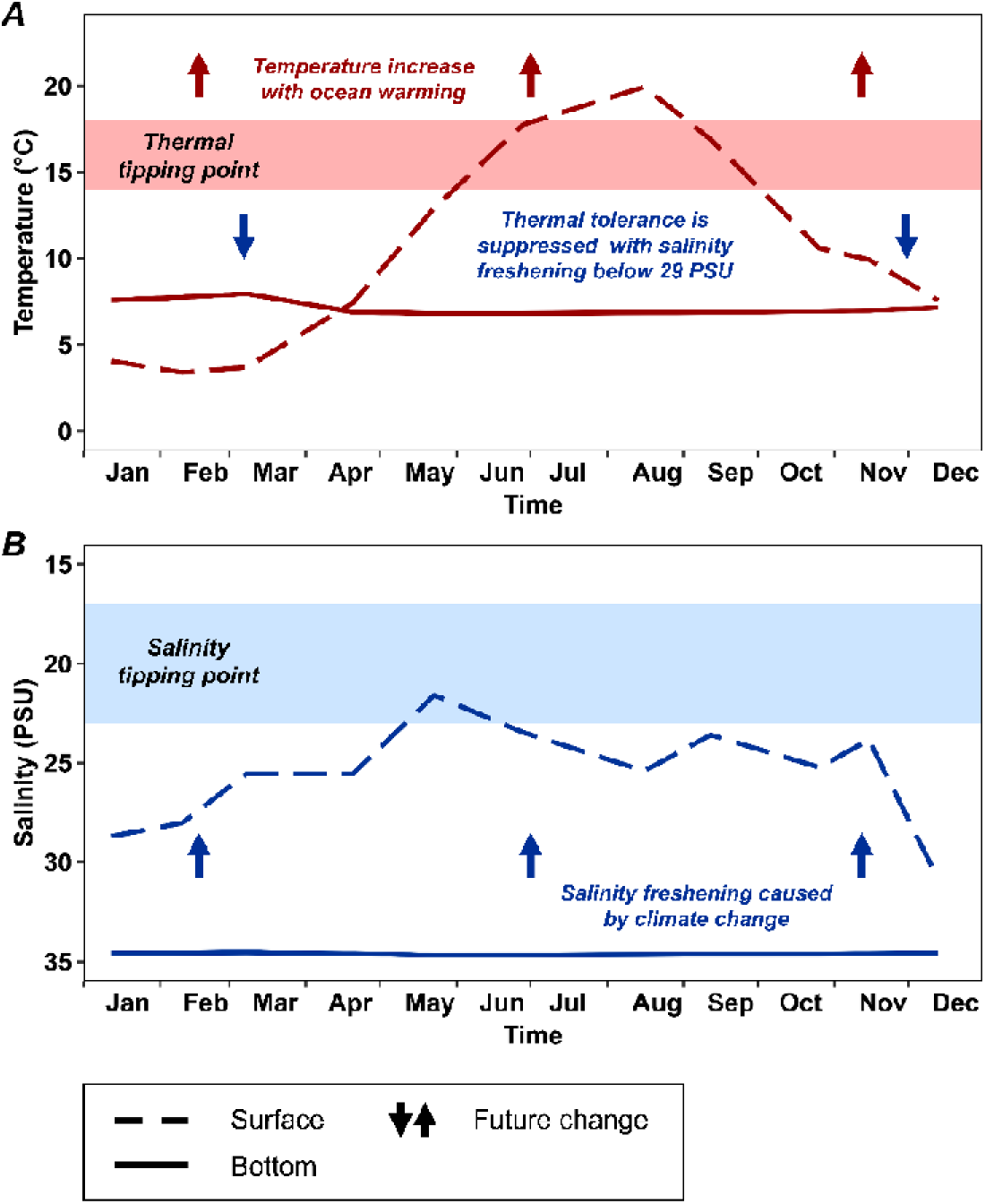
Temperature and salinity variations at the IM2 sampling station in the Oslofjord were recorded monthly in 2022. Temperature and salinity data are linearly interpolated over time between two measurement points. A) Temperature and B) salinity recorded at the surface (5 m below the surface, dashed line) and close to the bottom (198 m, 5 m above the bottom, full line).

We find that only salinity induced significant DNA damage. DNA damage increased below the tipping point for survival, at 23–17 PSU, and thus could be one of the processes driving mortalities. DNA damage occurs continuously in organisms due to endogenous factors, mainly through the production of reactive oxygen species (ROS) via metabolism [62–66]. Thus, organisms must continually repair DNA to ensure their survival. Indeed, DNA damage by altering translation-transcription affects all physiological processes, induces high energy costs via DNA repair, and can lead to cell death [62–66]. DNA damage is also triggered by exogenous factors [39,53,67,68], including SSF, which induced damage in fish, abalone, and shrimp, mainly due to increased ROS production during osmotic stress responses [69–72]. Temperature has also been reported to induce DNA damage in many species [73], although we observed no effect in *Calanus* spp.. It is important to note that DNA damage is not measurable in dead copepods due to rapid DNA degradation. Therefore, it cannot be excluded that it played a role in the warming-induced mortalities. In this case, the lack of damage in survivors could result from heat resistance or better DNA repair capacity. The disappearance of DNA damage with acclimation to stress can be observed in marine organisms [74–76].

The study of sublethal effects when stress exceeds tipping points is important because they can have lasting negative impacts on survivors, even after the stress is removed [77]. DNA damage can impact long-term fitness due to the energetic costs of DNA repair, aging, and mutations that can cause cancer and impair reproduction [62–66,78]. Therefore, exceeding the SSF tipping point at 23-17 PSU for a few days could have lasting effects on copepod fitness that could far exceed the short duration of our experiment. FMM determines the net amount of DNA damage, detected as single- and double-stranded DNA breaks and alkali labile sites, which result from both concomitant damage and repair [53,67]. Additional analyses could differentiate between DNA damage and repair processes (e.g., quantifying oxidised purines and pyrimidines, DNA methylation) [79,80]. Longer-term experiments could help understand how DNA damage and repair processes affect organisms’ fitness beyond tipping points.

The performance curves of survival as a function of heat accumulation (°C.d) provided further insight into the thermal tolerance of copepods. Importantly, we find that at a salinity of 29 PSU, heat accumulation does not affect survival until it exceeds a tipping point at 28°C.d, above which survival is affected. Heat accumulation corresponds to the time spent at a temperature above 8°C, the control temperature. Heat accumulations of 28 °C.d have only been exceeded in the incubators where copepods were exposed to temperatures above 14 °C (Supplementary Figure 2). Therefore, this supports the tipping point we observed for survival with the general Cox model between 14-18 °C. Studies on corals have already suggested the existence of a tipping point of heat accumulation above which survival drops [25,26]. Here, we first statistically determine the existence of such a tipping point. Reaching this tipping point for *Calanus* spp. could be equivalent to spending 28 days at 9°C or 2.33 days at 20°C.

Here, we did not test the influence of the ramping rate of temperature on the reaching of the survival tipping point of copepods. However, it is likely that reaching 28 °C.d in a short time would not have the same impact on survival as reaching this value over a longer time. For example, the influence of the ramping rate on influencing the CTmax of organisms has already been demonstrated [81]. Recovery phases after heat accumulation (periods of time when heat stress is removed) must also be taken into account. Previous studies on corals highlight two main cases in which prior heat accumulation 1) enhances or 2) reduces survival to a new MHW [25,27]. In copepods, prior heat accumulation negatively impacted the survival of *Tigriopus californicus* (Norman, 1869) exposed to a new MHW [28]. This suggests that, sometimes, the “heat accumulation debt” is not alleviated during a recovery period and that the tipping point is reached faster when the stress reappears. Further studies will need to determine whether different ramping rates and exposure cycles result in different heat accumulation tipping points.

Here, we find that SSF affects the thermal tipping point for survival in *Calanus* spp.. Previous study already showed that tipping point may arise earlier under the influence of an additional stressor [20,23]. Here, we find that exposure to additional stress causes the tipping point of a first stressor to disappear. This means that the phenotypic plasticity of organisms is no longer able to buffer the negative effects of stress on survival [17,37,38]. Indeed, SSF decreasing salinity from 29 to 23 PSU led to the immediate decrease of survival as soon as heat accumulates, with no tipping points. This decrease in survival is even steeper at 17 PSU. A recent study showed that decreases in CTmax and steepening of the slopes of thermal performance curves are frequent in marine species under SSF [24]. This is mainly due to the high energetic costs of osmoregulation via active ion transports, which compete with resources needed for thermoregulation [24,82,83]. Surprisingly, we find that salinity increasing from 29 to 32 PSU also causes the disappearance of the tipping point, even if the effect is lower than with SSF. Salinities of 29 to 32 PSU are present all year round below the pycnocline in the Oslofjord, where temperatures remain stable at 8 °C (Figure 5). Therefore, negative effects of increasing salinity on the resistance to warming are unlikely to occur in *Calanus* spp. habitats.

Climate change will narrow down the ecological niche of *Calanus spp.* populations. Indeed, OW and MHW will increase the duration of events when heat accumulates above 8°C in surface waters, thus accelerating the reaching of the tipping point (Figure 5). Concomitantly, while surface salinity is already below 29 PSU year-round in the Oslofjord (Figure 5), SSF will decrease it even further. Such low salinities will suppress the *Calanus* spp. thermal tipping point, i.e., the capacity of phenotypic plasticity to buffer negative effects of heat accumulation on survival. Deep waters represent a stable refuge where temperature and salinity remain stable around 8°C and 34.5 PSU year-round (Figure 5). However, OW and SSF will reduce the periods favourable for *Calanus* spp. to stay at the surface. *Calanus spp.* perform daily and seasonal vertical migrations to remain in optimal conditions for their biology [34,48]. Unfavourable surface conditions reduce their access to food that concentrates above the pycnocline, which impacts population dynamics [84]. Climate change is intensifying the stratification of the water column, deepening the pycnocline, and increasing salinity and temperature gradients [32]. This will further reduce the ecological niche available for *Calanus* spp., which could induce major changes in pelagic ecosystems.

## Conclusion

We show that the thermal tipping point of *Calanus* spp. at 14–18 °C is already reached in the surface waters of the Oslofjord during summer months. Decreasing salinities below 29 PSU already occurs in surface waters and will be further amplified by SSF. We found that such decreases in salinity suppress the ability (tipping point) of *Calanus* spp. to resist heat stress using phenotypic plasticity. Therefore, climate change will reduce the time that *Calanus* spp. can spend at the surface, reducing their access to food. This would affect the structure of pelagic ecosystems where *Calanus* spp. is the main link between primary producers and higher trophic organisms such as fishes [36]. Here, we show that the intensity and duration of exposure to warming determine the reaching of a tipping point of heat accumulation, beyond which survival is affected. We also demonstrate for the first time that exposure to a secondary stressor, SSF, can suppress tipping points, making the impact of a secondary stressor, OW, immediate on survival. We recommend increasing the complexity of experimental protocols for studying physiological tipping points. Progress could be made by studying the influence of ramping rates and exposure cycles (pulse v/s chronic exposures) on tipping points. When tipping points in phenotypic plasticity are reached, populations must adapt over the long term or risk extinction [12,13]. Rapid adaptation to ocean warming, over a few generations, has already been demonstrated in copepods [85]. Therefore, studies determining the evolution of physiological tipping points over multiple generations are the next step in projecting the future of species. Such studies are needed in a world where tipping points are increasingly crossed by environmental changes [21,86].

## Data availability

All data presented in this study and the R scripts used to analyse them are available in a data repository (10.6084/m9.figshare.29433440). Complementary information is available from the corresponding authors on reasonable request.

## Authors’ contributions

M.L.: conceptualization, data curation, formal analysis, investigation, methodology, validation, writing - original draft; A.E.T.S.: conceptualization, data curation, formal analysis, investigation, methodology, validation, writing - original draft; H.R.: investigation, methodology, validation; H.S.K.: investigation, methodology, validation; K.B.: investigation, methodology, validation; K.V.D.: conceptualization, investigation, methodology, validation, supervision, funding acquisition, project administration, writing - review and editing. All authors gave final approval for publication and agreed to be held accountable for the work performed therein.

## Acknowledgments

The authors thank research groups at the University of Oslo and the Scottish Association for Marine Sciences for assistance with field sampling and DNA damage analyses.

