## Supplementary Figure 1 for "Sea surface freshening can suppress the thermal tipping point of marine copepods"

Running head: Salinity effects on copepod thermal tipping point

^†^ Co-first authors

### Figures

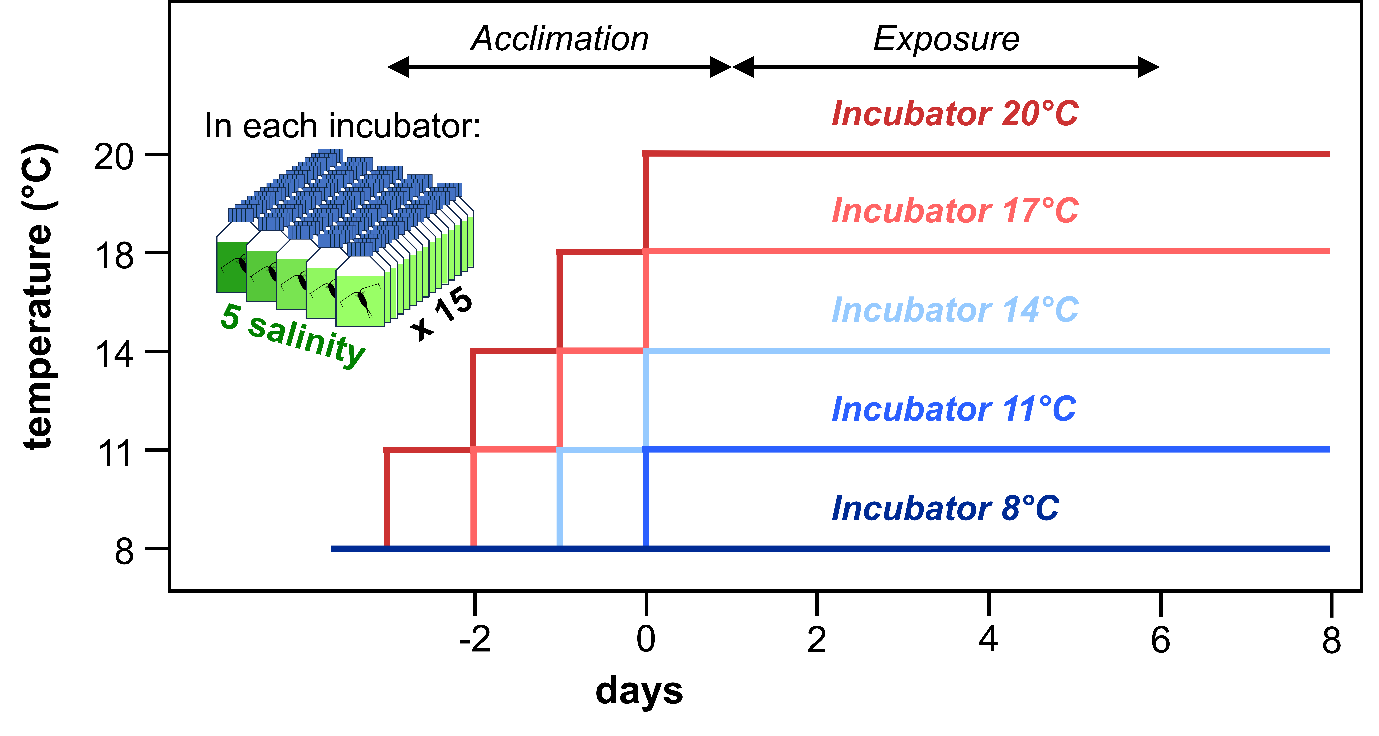

**Supplementary Figure 1.** Experimental design used to determine the temperature tolerance of *Calanus* spp. from the Oslofjord across 5 salinity conditions. Variations in temperature over time are presented for the 5 incubators, each containing 75 copepodite V *Calanus* spp., i.e., 15 per salinity conditions. The salinity conditions were 32, 29, 23, 17, and 10 PSU. The first three days corresponded to the gradual increase in temperature in the incubators. The exposure then lasted 8 days for all temperature conditions.

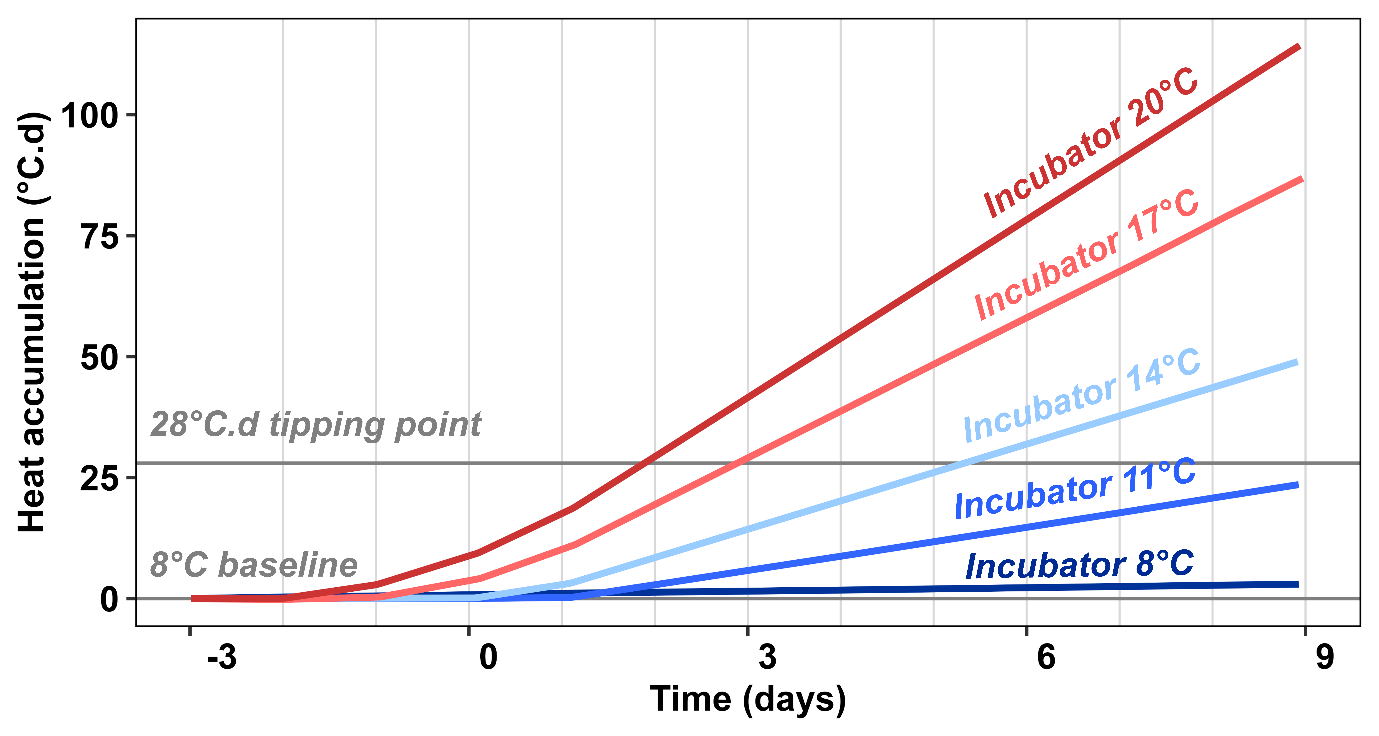

**Supplementary Figure 2.** Variation of heat accumulation (°C.d), measured in each of the 5 incubators, over time. Each incubator is represented with a different colour. Negative values for time on the x-axis indicate the days before the designated experimental temperature condition is reached, i.e., the “acclimatisation” period. The exposure period is indicated by positive values. The light grey vertical lines represent the times when survival was checked in the incubators (once per day). The two horizontal lines represent the 8 °C control temperature (0 °C.d) and the heat accumulation tipping point (28 °C.d) at 29 PSU, respectively.

### Tables

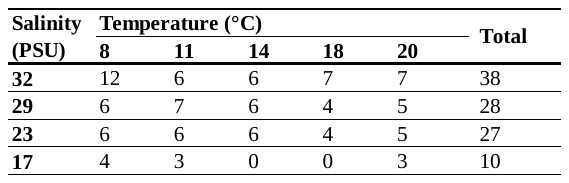
**Supplementary Table 1.** Number of copepods analysed for DNA damage for each temperature and salinity condition.
